# Study of flow effects on temperature-controlled radio-frequency ablation using phantom experiments and forward simulations

**DOI:** 10.1101/2020.06.26.172957

**Authors:** Teresa Nolte, Nikhil Vaidya, Marco Baragona, Aaldert Elevelt, Valentina Lavezzo, Ralph Maessen, Volkmar Schulz, Karen Veroy

**Affiliations:** Department of Physics of Molecular Imaging systems, Institute for Experimental Molecular Imaging, RWTH Aachen University, Forckenbeckstr. 55, 52074 Aachen, Germany; Aachen Institute for Advanced Study in Computational Engineering Science, Schinkelstrasse 2, 52062 Aachen, Germany; Philips Research Europe, High Tech Campus, 5656 AE Eindhoven, The Netherlands; Hyperion Hybrid Imaging Systems GmbH, Pauwelsstr. 19, 52074 Aachen, Germany; Physics Institute III B, RWTH Aachen University, Aachen, Germany; Fraunhofer Institute for Digital Medicine MEVIS, Aachen, Germany; Center for Analysis, Scientific Computing, and Applications, Eindhoven University of Technology, 5600 MB Eindhoven, Netherlands

**Keywords:** temperature controlled radiofrequency ablation, treatment planning, forward simulation, flow effects

## Abstract

**Purpose:** Blood perfusion is known to add variability to hepatic radiofrequency ablation (RFA) treatment outcomes. Simulation-assisted treatment planning taking into account blood perfusion may solve this problem in the future. Hence, this study aims to study perfusion effects on RFA in a controlled environment and to compare the outcome to a prediction made using finite volume simulations.

**Methods:** Ablation zones were induced in tissue-mimicking, thermochromic ablation phantoms with a single flow channel, using a RF generator with needle temperature controlled power delivery and a monopolar needle electrode. Channel radius and saline flow rate were varied and the impact of saline flow on the ablated cross-sectional area, on a potential occurrence of directional effects as well as on the delivered generator power input was studied. Finite-volume simulations reproducing the experimental geometry, flow conditions and generator power input were conducted in a second step and compared to the experimental ablation outcomes.

**Results:** Vessels of different radii affected the ablation result in different manners. For the channel radius of 0.275 mm both the ablated area and energy input reduced with increasing flow rate. For radius 0.9 mm the ablated area reduced with increasing flow rate but the energy input increased. An increasing area and energy input were observed towards larger flow rates for the channel radius of 2.3 mm. Directional effects, i.e., shrinking of the lesion upstream of the needle and an extension thereof downstream, were observed only for the smallest channel radius. The simulations qualitatively confirmed these observations. When using the simulations to make a prediction of ablation outcomes with flow, the mean absolute error between experimental and predicted ablation outcomes was reduced from 23% to 12% as compared to neglecting flow effects.

**Conclusion:** Simulations can improve the prediction of RFA ablation regions in the presence of various blood flow effects. Our findings therefore underline the potential of simulation-assisted, patient-individual RFA treatment planning and guidance for the prediction of RFA outcomes in the presence of blood flow.

**Additional comments:**

-Teresa Nolte and Nikhil Vaidya contributed equally to this work.
-Volkmar Schulz and Karen Veroy contributed equally to this work.
-A single reference experiment, i.e., not using a flow channel, and the image in the upper left corner of Figure 4 were included into a publication submitted to Int. J. Hyperthermia for model validation purposes.

## 1 Introduction

Percutaneous thermal ablation techniques can treat both primary and secondary focal liver tumors in a minimally invasive manner [1]. However, radiofrequency ablation (RFA) outcomes in the liver cannot be easily predicted due to the cooling effect of blood flow, the complexity of the vessel architecture and variable liver properties [2] [3]. Therefore, RFA treatments are commonly evaluated by a contrast-enhanced CT scan [4], permitting the clinician to decide if the tumor region (including a safety margin) has been ablated. Compared to other clinically established ablation techniques such as microwave ablation, RFA is preferred for treatments near adjacent sensitive structures [5]. Treatment planning for RFA involving numerical simulations based on the patient-specific geometry aims to suggest optimal treatment parameters to the clinician in order to ensure complete ablations while limiting damage to adjacent structures [6]. This promises to ease the burden for the patient by reducing tumor recurrence. The ablation zone is affected by the distributed perfusion of small blood vessels, and also by large vessels which cause localized heat transport [7]. Depending on the vessel radius and the blood flow rate there is a size reduction and deformation of the ablation zone and/or stretching of the thermal lesion in the flow direction [7] [8]. The latter will be referred to as directional effect throughout this paper. Consequently, a more realistic treatment outcome can be predicted by including perfusion effects into treatment planning.

Modeling blood-flow effects is complicated due to the diversity in blood-vessel diameters and topologies. A study by Pennes [9] used a model in which small, densely packed blood vessels were modelled by a volumetric heat sink term. Work by Nakayama and Kuwahara [10] derived a more realistic model for the heat sink effect of small blood vessels. This two-equation model treats blood temperature and tissue temperature as coupled variables. To our knowledge, the directional effect of individual blood vessels has not been studied in detail. For example, it is unclear where to place the radius cut-off to distinguish between large and small vessels. We have, in the past, performed a simulation study on the directional effect of blood vessels in RFA [11]. While assuming a dependence of flow rate on the vessel radius as suggested by Murray’s law [12], directional effects were found to occur below vessel radii of 0.5 mm and, if considering blood coagulation effects, above 0.4 mm for the blood vessel parameters considered. Kim et. al. [13] have reported that a safety margin greater than 3 mm induced in clinical treatment appears to be associated a lower rate of local tumour progression. It may be important to determine whether any directional effects interfere with this margin with respect to treatment success.

**Table 1.**
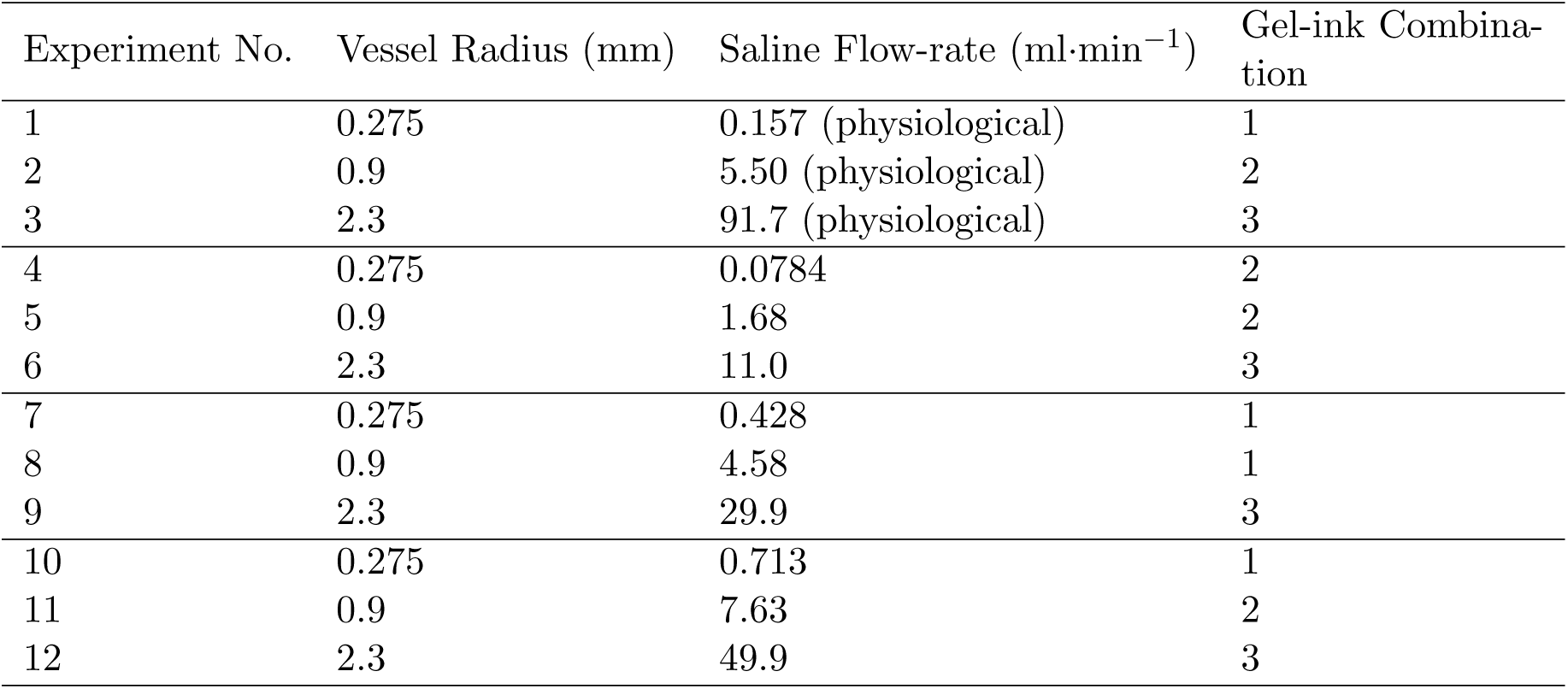
Table showing vessel radius, saline flow rates and gel-ink combinations used for the experiments performed. Experiments 1-3 used the physiological flow rates for a given vessel radius according to Murray’s Law [12].

The impact of blood perfusion on RFA outcomes has been studied experimentally. Next to the retrospective evaluation of clinical RFA treatments [2][14], many studies have been conducted in ex-vivo liver setups. Using an impedance-controlled radiofrequency (RF) system with constant power input and LeVeen electrodes in machine-perfused bovine livers, Bitsch et. al [7] observed both size reductions and deformations of ablation zones under perfusion in comparison to non-perfused controls. Pillai et al. [15] evaluated monopolar RFA electrodes in the vicinity of major hepatic veins in a perfused calf liver using a temperature-controlled RF generator. They found reduced volume and mass of the ablated lesions, along with longer ablation times needed for the treatment. The study of Poch et. al. [16] examined flow through a perfused glass tube within ex-vivo tissue using multipolar electrodes and an impedance-controlled RF generator with automatically modulated power. Vascular cooling resulted in incomplete ablation at different vessel-to-applicator distances and changes in shape. Another study [17] compares ablation results in a perfused glass vessel setup to a finite element model. A monopolar ablation needle with a cooled tip was used in conjunction with an impedance-controlled RF generator and constant power settings. Blood flow did not affect the lesion diameter for large distances between electrode and vessel, although shape deformations seemed to occur. González-Suárez et. al. [18] presented an in vivo study for an RFA assisted liver resection device under ultrasound guidance and found agreement between experimental and simulated lesion shapes. The simulations were subsequently used to predict whether the heat-sink effect can itself protect the portal vein, thereby pointing out the potential of treatment planning approaches in an in-vivo setting.

The objective of the present study is to demonstrate that treatment planning for RFA interventions in the presence of perfusion can improve the prediction of ablation outcomes. We use a temperature-controlled RF generator and ablation phantoms with a single flow channel to study ablation outcomes for defined vessel radii and saline flow rates and compare the results to simulations (using the finite volume method). Thereby, we evaluate how well the simulations predict the ablation outcomes under known flow conditions. We assess the resulting ablation zone areas as well as the occurrence of directional effects. Moreover, the impact of flow on the generator feedback algorithm is studied. As compared to in- and ex-vivo liver experiments, the controlled environment and simplified geometry established in this study permits to investigate the effects of flow on ablation results in a controlled manner. Potential implications of the study towards the in-vivo treatment situation will be discussed.

## 2 Methods

### 2.1 Experimental Setup

#### 2.1.1 Fabrication of ablation phantoms

Tissue-mimicking thermochromic phantoms consist of a polyacrylamide gel formulation, to which a thermochromic ink as well as sodium chloride are added. The ink changes color irreversibly from white to magenta above a threshold temperature, while sodium chloride mimics the electrical conductivity of liver tissue. Discoloring has been described as proportional to the maximum achieved temperature value [19][20]. For phantom preparation, 671 g of deionized water were mixed with 175 g of a 40% (Bis-)Acrylamide 19:1 solution. Fifty gram of irreversible change ink concentrate and 7 g of sodium chloride (Sigma Aldrich, USA) were added to the solution and mixed under continuous stirring. 1.4 g of the catalyst N,N,N0,N0- tetramethylethylenediamine was added before 1.4 g of ammonium persulfate (both reagents purchased from Sigma Adrich, USA), which initialized polymerization. The mixture was poured in a cube-shaped container and allowed to polymerize at room temperature. The different gel-ink combinations used are shown in table 2. The gel-ink combination used for each of the experiments is shown in table 1. The Kromagen Magenta MB60-NH and 60-NH inks will be referred to as ink I and ink II respectively.

**Table 2.**
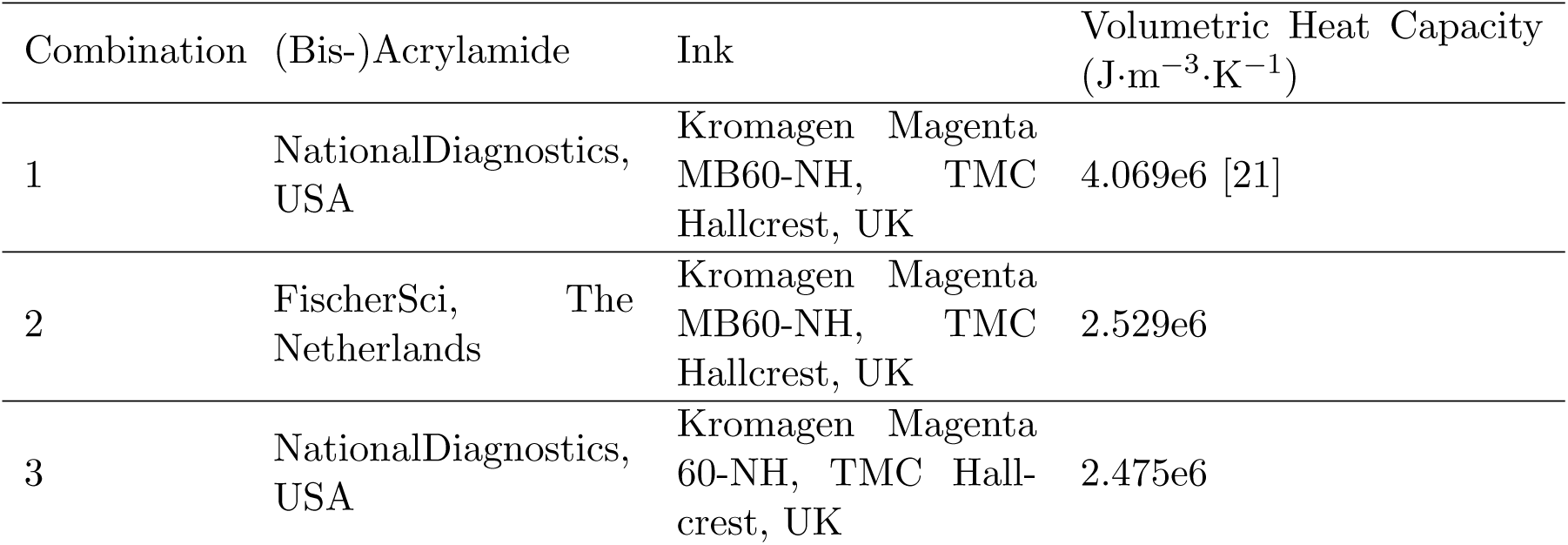
Different gel-ink combinations used in the study and their respective volumetric heat capacities.

The custom-fabricated sample container (Figure 1) allowed to insert rods of different radii through the center of the cube before pouring in the liquid phantom material. The inner dimensions of the sample container were 10 cm x 10 cm x 10 cm. After polymerization, the rods were removed without damaging the now solidified phantom, leaving a flow channel of known position and dimensions. Moreover, the cuboid container provided an insertion channel for the monopolar RFA needle to position it parallel to the flow channel at a wall-to-wall distance of 3 mm. The 10 x 10 cm^2^ sized ground pad (Angiodynamics, New York, USA) was located on the bottom face of the sample container. Figure 2a shows a coronal cross-section through needle and flow-channel as well as a 3D view of the geometry.

**Fig. 1.**
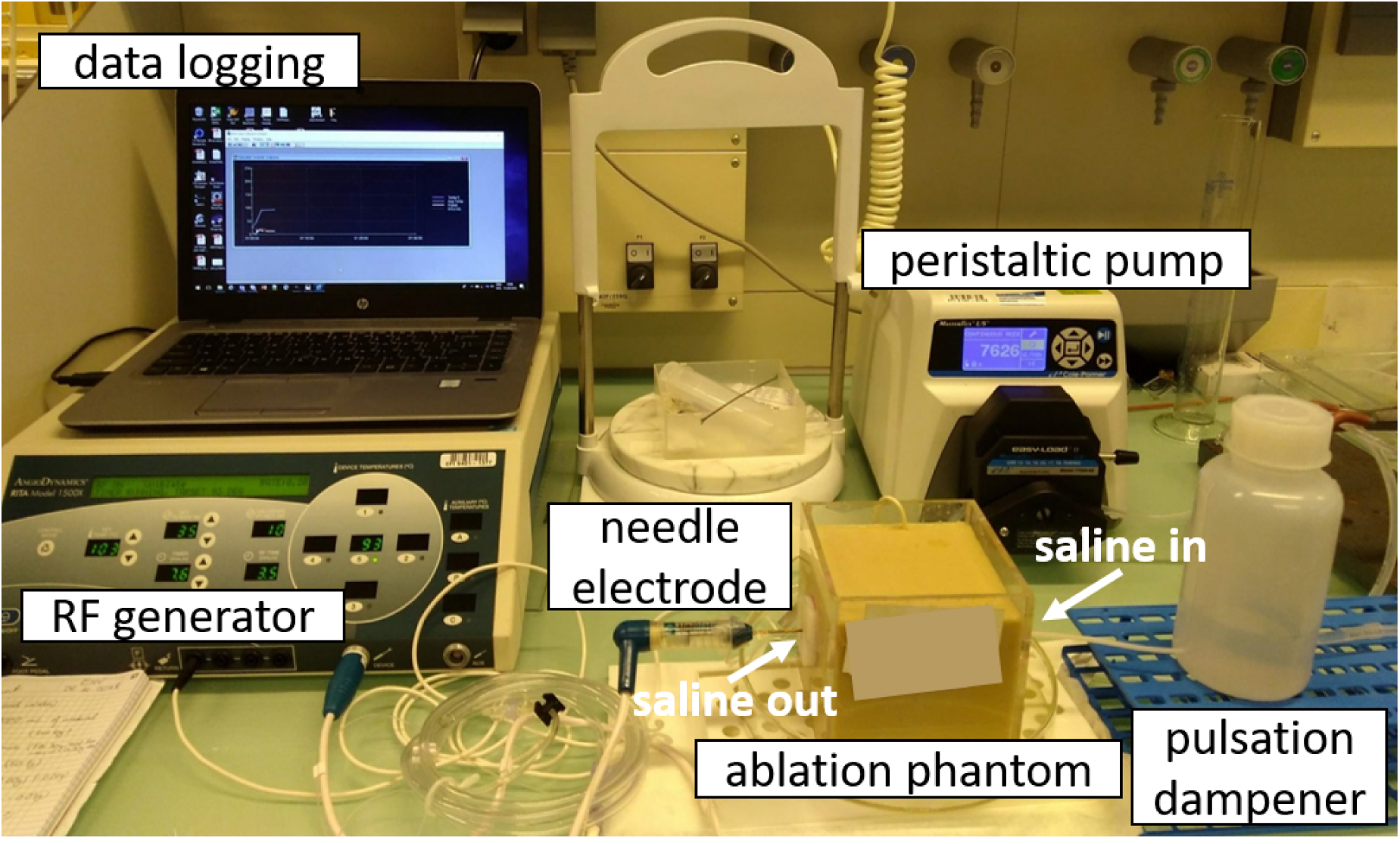
Photograph of experimental setup showing the box containing: (i) ablation phantom with a flow channel; (ii) the RF generator with data logging and needle electrode installed within the phantom, and (iii) flow circuitry including a peristaltic pump, pulsation dampener and tubing. Saline outflow from the phantom occurred on the needle side.

**Fig. 2.**
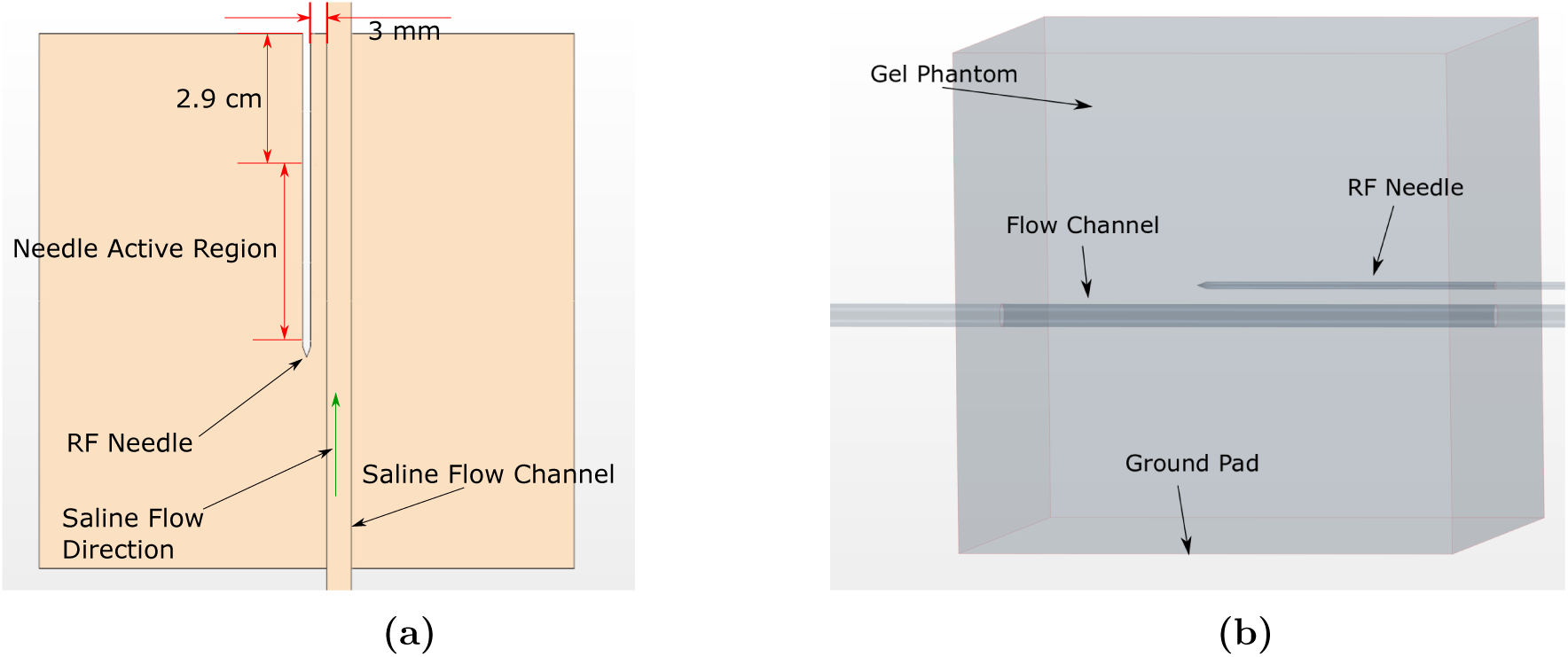
**(a)** Top view of geometry showing plane containing the ablation needle and saline flow channel axes; **(b)** 3D view of phantom cube, ablation needle, and saline flow channel

#### 2.1.2 RFA experiments including flow

Table 1 lists the RFA experiments performed. Three flow channel radii (0.275 mm, 0.9 mm, and 2.3 mm) and four distinct flow rates per channel were considered. This setup made it possible to independently study the influence of channel radius and flow rate on the directional effect of the saline flow. The flow rates included a “physiological” flow rate for each channel radius, assumed to be given by Murray’s Law [12]. RFA experiments were carried out using a RITA 1500X RF generator and a Uniblate ablation needle (both by Angiodynamics, NewYork, USA) with an active electrode length set to 3 cm. Temperature setpoint, ablation time, and maximum power were 103°C, 10 min, and 35 W, respectively. Saline solution (0.1% (w/w) in dionized water) was prepared and flow through the channel was established using either a Multi-Phaser™ NE-1000 syringe pump (ProSense, Oosterhout, the Netherlands) or a Masterflex L/S peristaltic pump (Cole-Parmer, Wertheim, Germany). For experiments with the latter one, a custom-made pulsation dampener was inserted into the flow circuit to ensure uniform flow. Saline flow was turned on prior to ablation and stopped after the generator’s cool-down cycle. During each RFA experiment, the delivered power as well as the temperature at the needle tip were logged from the generator every second. Reference RFA experiments were also conducted, omitting the flow channel during phantom preparation and conducting the RFA experiment with identical settings otherwise. After each experiment, the ablation needle was removed and the phantom material was cut in half along the central plane passing through both needle canal and flow channel. The resulting discolored zone was photographed with a custom-made camera setup including a CCD camera (IDS Imaging Development Systems, Obersulm, Germany) and reproducible lighting conditions.

### 2.2 Analysis of ablation zones

Photographs of the discolored ablation zones were analyzed using the open source software GIMP. For both ink I and ink II, the photographs mentioned above were transformed to gray values and the discolored zone was thresholded. For phantoms prepared with ink I, preliminary calibration experiments showed that full discoloring was achieved after heating the material above 70°C. For phantoms prepared with ink II, discoloring was mostly, but not yet fully completed at the lower temperature of 65°C. To still assess the *T* > 70°C zone, we used the gray value of the 70°C calibration experiment for thresholding. The area of the *T* > 70°C zone was computed using GIMP’s histogram tool. The simulated *T* > 70°C zone was overlaid with the photographs by aligning needle channel, flow channel and outer border of the phantom cube.

### 2.3 Simulation Setup

#### 2.3.1 Simulation Geometry

The simulation geometry consisted of three subdomains: (i) thermochromic gel phantom, (ii) ablation needle, and (iii) saline flow channel. The Uniblate ablation needle was modelled with exact outer dimensions, while average thermal and electrical properties were used in the interior. Figure 3 shows a close-up of the ablation needle subdomain. The needle active region is represented as a separate boundary surface which allows electric current to pass, while the two insulated surfaces do not. The top surface of the needle is used to apply the input voltage which induces tissue heating.

**Fig. 3.**
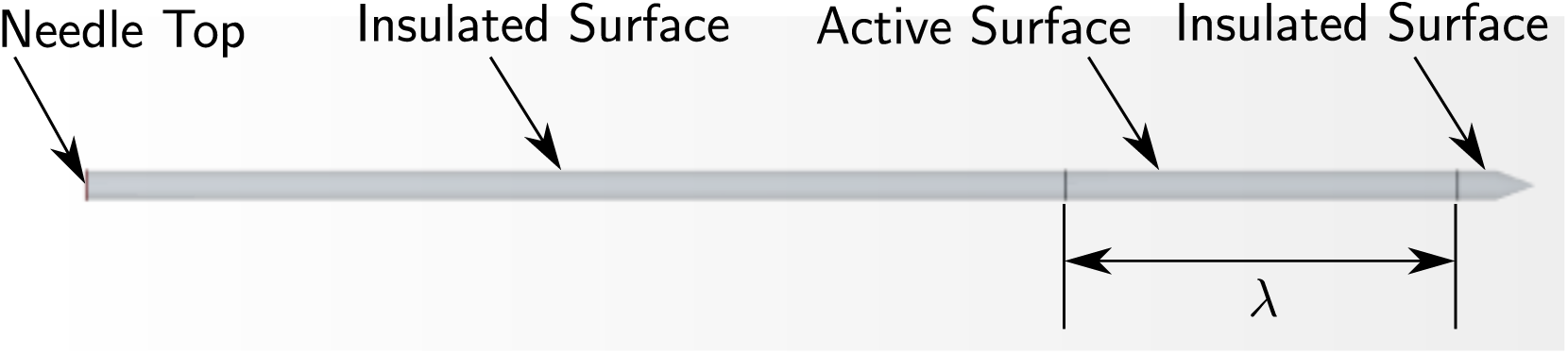
Enlarged view of the ablation needle geometry

#### 2.3.2 Mathematical Model

In this work, heat transfer in solid thermochromic phantom was described by the diffusion equation, while that in saline was described by advection-diffusion. The governing equations for gel phantom (*p*), saline (*s*), and ablation needle (*n*) subdomains are respectively given by 

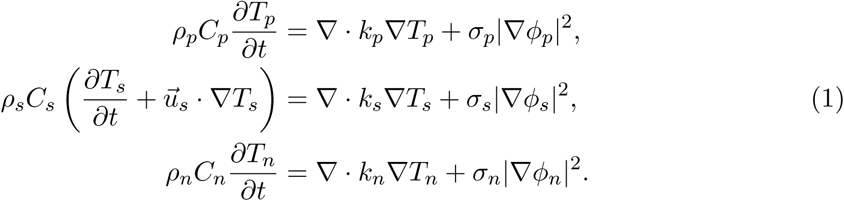

where *ρ*_*i*_, *C*_*i*_, *T*_*i*_, *k*_*i*_, *σ*_*i*_, and *ϕ*_*i*_ are the density, specific heat, temperature, thermal conductivity, electrical conductivity, and electric potential, respectively, and *i* ∈ {*p, s, n*}. Similarly, the expressions *σ*_*i*_|∇*ϕ*_*i*_|^2^ are the RF heat source densities in the three subdomains. The saline velocity 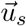 is assumed to have a laminar and steady profile. The temperature and heat-flux are continuous across all subdomain boundaries. The electric potential in all subdomains is governed by the quasi-static approximation 

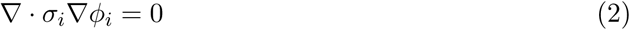

where *i* ∈ {*p, s, n*}. A time-dependent voltage 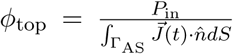 is applied at the top boundary of the ablation needle, where *λ* is the active length of the ablation needle, Γ_AS_ is the active surface of the needle, and 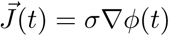 is the electric current density. Functions *ϕ* and 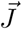 are continuous across the vessel-tissue boundary and the active surface of the needle. The input power, *P*_in_, is taken from the RF generator logging data. The bottom surface of the phantom was at (ground pad) 0 V, while all other boundaries were electrically insulated. The saline inflow and outflow boundaries were kept at room temperature. All external boundaries of the phantom use the thermal boundary condition 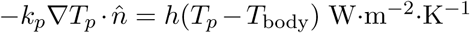. The needle surface exposed to the air also had a convection boundary condition. The values of *h* for these boundaries are given Table 3. The initial temperature was 21°C.

**Table 3.** Convection coefficient values for external boundaries.

| Boundary | Convection Coefficient<br>$h$ ($\text{W}\cdot\text{m}^{-2}\cdot\text{K}^{-1}$) |
| --- | --- |
| Top Phantom Face (air inter-<br>face) | 12 |
| Bottom Phantom Face (rest-<br>ing on table) | 9 |
| Phantom Side Faces (parallel<br>to ablation needle and flow-channel) | 12 |
| Phantom Side Faces (perpen-<br>dicular to ablation needle and<br>flow-channel) | 10 |
| ablation needle - Air Interface | 12 |

The density, specific heat capacity, and thermal conductivity of the thermochromic gel phantom for the gel-ink combination 1 were taken from [21]. The gel-ink combinations 2 and 3 have different heat capacities, which were estimated from their cooling behavior, given in table 2. The complex dielectric properties of the gel were measured and a conductivity value of 1 S·m^−1^ was obtained at the generator frequency (460 MHz). The saline density *ρ*_*s*_ = 970 kg·m^−3^ and specific heat capacity *C*_*s*_ = 4200 J·K^−1^·m^−3^ were taken from [22] [23], using the temperature dependent expression. Median values of density and specific heat capacity were chosen from the temperature range of interest. The average material properties used for the needle were: *ρ*_*n*_ = 1 kg·m^−3^, *C*_*n*_ = 1000 J·kg^−1^·K^−1^, *k*_*n*_ = 0.02 W·m^−1^·K^−1^, and *σ*_*n*_ = 1.0*e*7 S·m^−1^.

## 3 Results

### 3.1 Influence of flow conditions on ablated areas

Figure 4 gives an overview of all ablation results. Below the corresponding reference experiment without flow channel, all experiments with saline flow are depicted. The experimental parameters (flow channel radius, flow rate) are indicated below the ablation zones. Around the needle, the magenta-colored zone is visible, indicating where tissue temperatures over 70°C or 65°C were reached for ink I or II, respectively. The experimental *T* > 70°C zones, obtained by gray-value based thresholding, are marked in blue, whereas the simulated *T* > 70°C zones are marked in black. Without flow, the ablated areas are symmetric with respect to the needle. For a radius of 0.275 mm and the two smallest flow rates of 0.078 ml·min^−1^ and 0.157 ml·min^−1^, the *T* > 70°C zones are distorted along the saline flow direction. These directional effects will be further discussed in section 3.3. For larger flow rates, the 70°C zones do not cover the flow channel region anymore, which reveals heat transport by saline away from the ablation site. The simulated *T* > 70°C zones show a close correspondence with the discolored areas in general. However, in four experiments the experimentally obtained 70°C zone area unexpectedly exceeds the simulated area towards the top end of the needle, i.e., for experiments (1), (5), (7) and (10). This feature will be referred to as the ‘bump’ structure in the following.

**Fig. 4.**
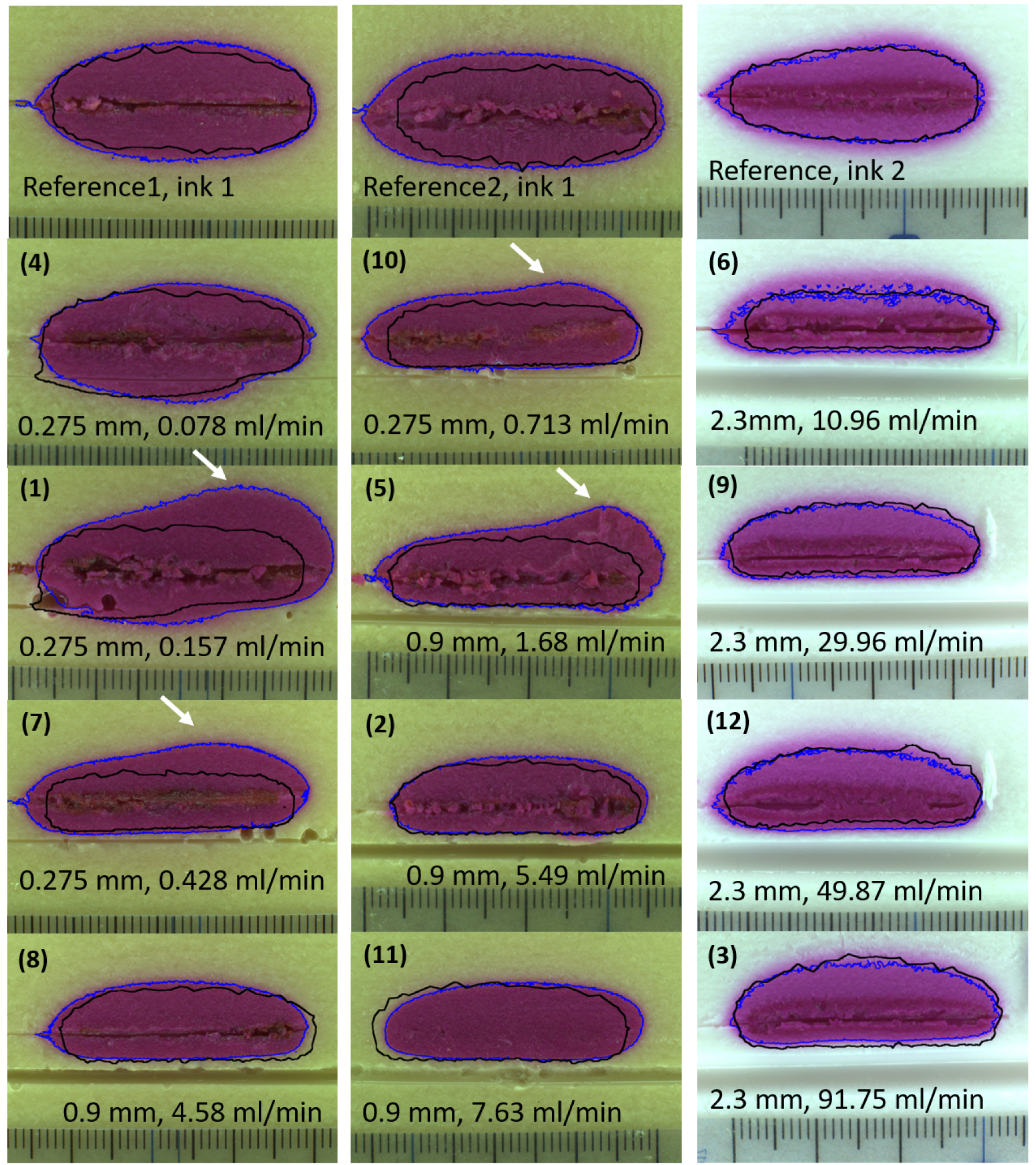
Experimentally obtained ablation zones for all RFA experiments and the corresponding reference experiments. The experimental and simulated 70°C temperature contours are marked by a blue and a black line, respectively. The ablation needle was inserted from the left side. Saline flow was established from right to left through the flow channel located below the ablation area. Radius and flow rate for each experiment were as specified within the figure below the ablation zones. In four experiments, the experimental ablation zone shows an unexpected ‘bump’ structure, which we marked by white arrows. The numbering of the experiments refers to Table 1.

Figure 5 shows the experimental and simulated normalized ablated areas, determined by the *T* > 70°C zone, for the different flow experiments. Normalization was performed with respect to the experimental or simulated area of the corresponding reference experiment. For the smallest radius of 0.275 mm, both normalized areas are found to decrease with flow rate apart from one outlier. In contrast, both simulated and experimental ablated area increase towards higher flow rates for the largest channel radius of 2.3 mm despite the increased heat transport away from the ablation site. For the intermediate channel radius of 0.9 mm, the experimental ablation area slightly decreases with flow rate, while the simulated ablation area slightly increases.

**Fig. 5.**
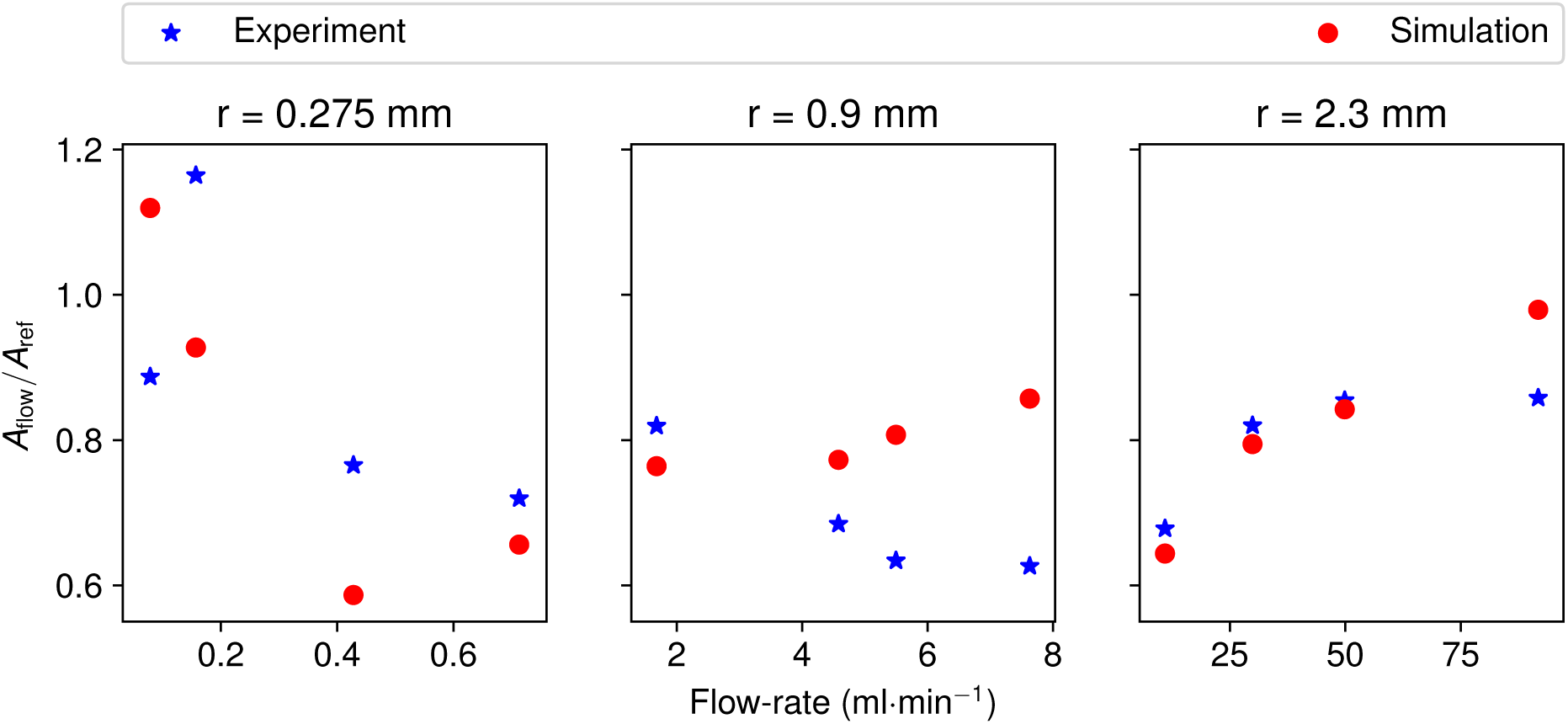
Experimental and simulated ablation zone areas of the flow experiments, normalized to the corresponding reference experiments, for the different investigated flow channel radii.

### 3.2 Comparison between simulation and experiment

The experimental and simulated T > 70°C zone areas are compared by means of the correlation plot in Figure 6 to assess if the simulations are able to predict the RFA treatment outcome in the presence of different flow conditions. The plot shows that the simulated *T* >70°*C* zones are close to the experimental ones. Some simulated zone areas underestimate the experimental ones, among which are the above-described ones exhibiting the ‘bump’ structure towards the top end of the ablation zone, cf. Figure 4. The deviation of our experimentally obtained normalized ablation areas, *a*_exp_ = *A*_flow, exp_*/A*_ref, exp_, from a predicted ablation outcome, *a*_pred_, was further quantified by mean absolute error (MAE) calculations, defined as 

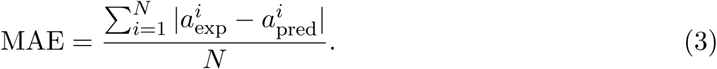

**Fig. 6.**
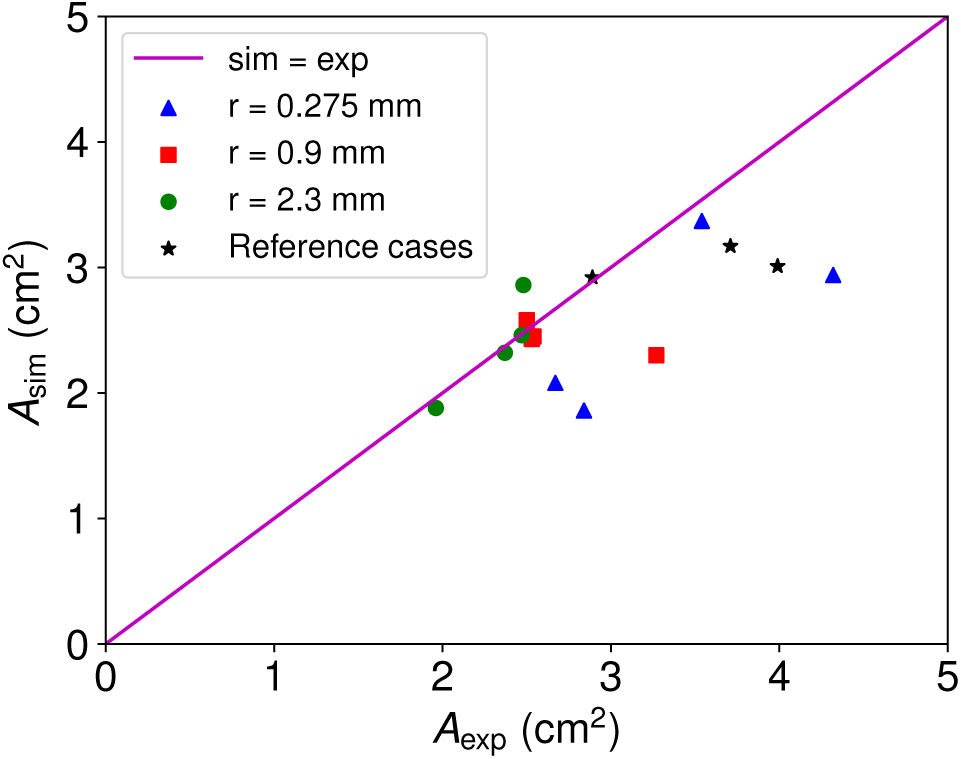
Correlation plot comparing the experimental and simulated T> 70°C zone areas *A*_exp_ and *A*_sim_. The simulations underestimate the experimental areas for the smallest channel radius. These are the experiments with the ‘bump’ structure in the *T* > 70°C zone.

Here, *N* = 12 is the number of experiments including flow channels. When using the simulated normalized areas to predict the ablation outcomes, i.e., setting *a*_pred_ = *a*_sim_, thereby quantifying the mean difference between the blue and red data points in Figure 5(a)-(c), MAE=0.12 is obtained. When neglecting flow effects from the prediction and assuming the outcome equals the outcome of the reference experiment without any flow, i.e., when setting *a*_pred_ = 1, a two-fold higher MAE=0.23 is obtained. This comparison clearly demonstrates that simulations can improve the prediction of treatment outcome in the presence of flow. It should be noted that a direct comparison of the experimental results to the vendor prediction was unfortunately not possible, as the electrode manual does not comment on the extent of the *T* > 70° zones.

### 3.3 Directional Effects

The directional effect of saline flow can be seen as a stretching of the *T* > 70°C region along the flow direction. In Figure 4 this phenomenon is visible only for experiments 1 and 4. According to our simulation experience, the thermal lesion in liver tissue typically coincides with a boundary temperature of around 52°C. The study by Huang [8] showed that lower temperature contours are stretched more by blood flow, so the *T* > 52°C zones were also obtained from the simulations. The simulation results for the 52°C contour show directional effects in experiments 1, 4, and 7 from Table 1. Figures 7a through 7e show cross-sections of the T > 52°C zones obtained in these experiments. The red outline shows the flow-affected lesion, while the blue outline shows the reference lesion. In experiments 1, 4, and 7 there is a distinct “tail” on the *T* > 52°C zone which stretches in the flow direction. For the channel radius of 0.275 mm and flow-rate 0.0784 ml·min^−1^, a small directional effect occurs in the *T* > 52°C zone. The directional effect is larger when the flow-rate rises to 0.157 ml·min^−1^. For the flow-rate of 0.428 ml·min^−1^, the directional effect decreases. Finally, for the flow-rate of 713 ml·min^−1^, the directional effect is negligible. The tail-structure due to the blood flow was not seen for any other channel radius. In all the figures, the vessel-affected and reference lesions differ only on the vessel-side of the ablation needle.

**Fig. 7.**
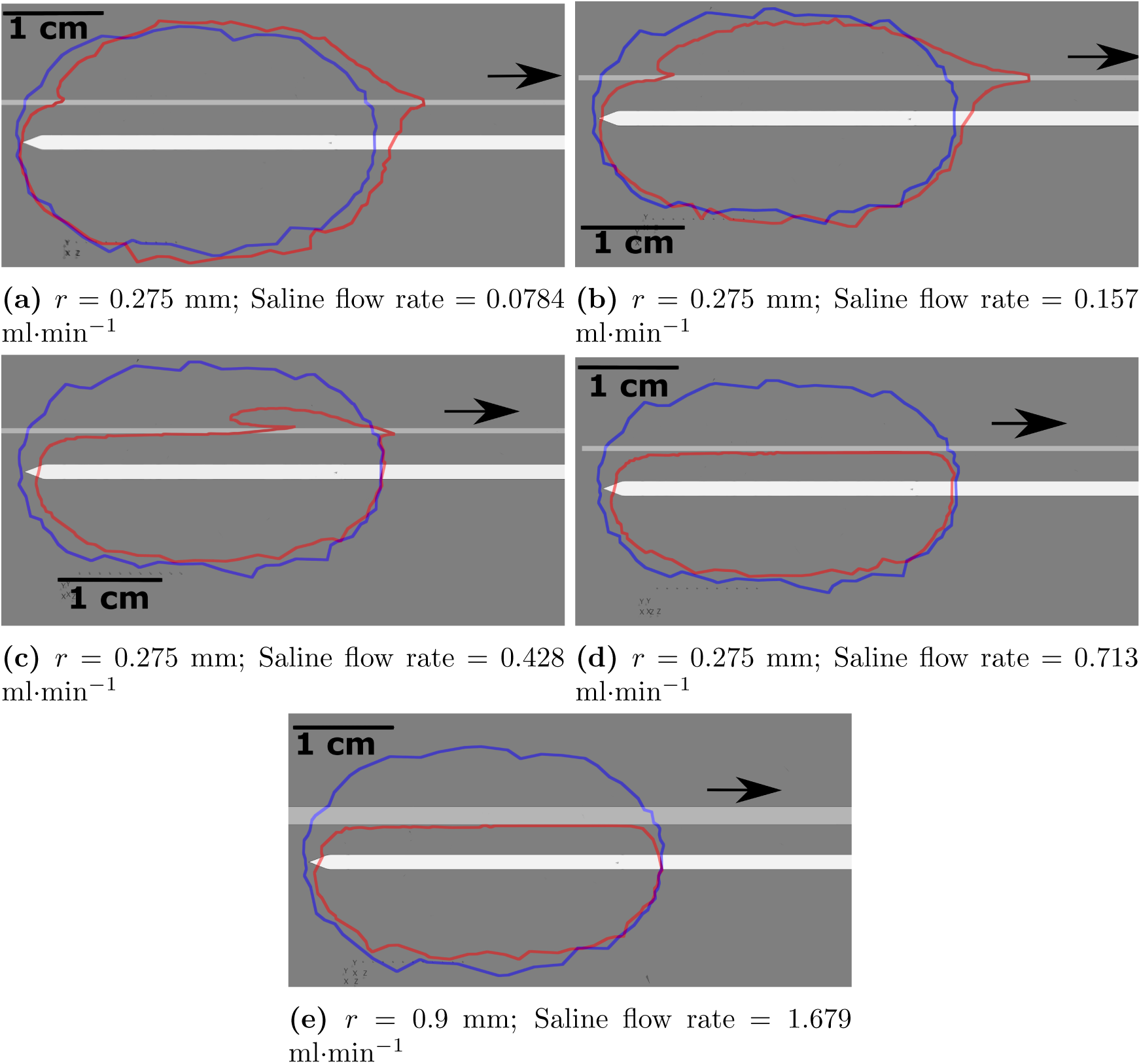
The reference *T* > 52°C zone (blue) and the flow-affected *T* > 52°C zone (red) for channel radius 0.275 mm with increasing saline flow-rate (subfigures (a) to (d)) and for channel radius 0.9 mm with the lowest flow-rate of 1.679 ml·min^−1^ (e). The black arrow indicates the saline flow direction

Figure 7b shows the longest tail structure in the thermal lesion of the different cases considered in this work. Also, the flow-affected *T* > 52°C zone deviates from the reference zone mostly near the entry and exit points of the flow. Near the inflow end, there is a shrinking of the *T* > 52°C of the order of 4 mm. Similarly, near the outflow end, there is tail region of 9 mm in length.

### 3.4 Analysis of generator power input

A temperature-controlled RF generator was used in this study. Hence, the input power over time is influenced by both experimental conditions (channel radius, saline flow-rate) and generator settings (temperature setpoint, maximum power, ablation time). From the logged generator power curves *P* (*t*), the total energy input was calculated as *E* = *P* (*t*)*dt* for all ablation experiments and normalized to the energy input of the corresponding reference experiment. An example for a power curve *P* (*t*) is shown in Figure 8(a). Oscillations in the curves probably originate from the generator feedback-loop, as the device constantly adjusts the power to follow a prescribed temperature curve which it monitors at the tip of the ablation needle [24]. According to subfigures 8(b) to (d), for the larger radii, i.e., 2.3 mm and 0.9 mm, the energy input increases with flow speed, whereas a decreasing energy input is observed for the smallest radius.

**Fig. 8.**
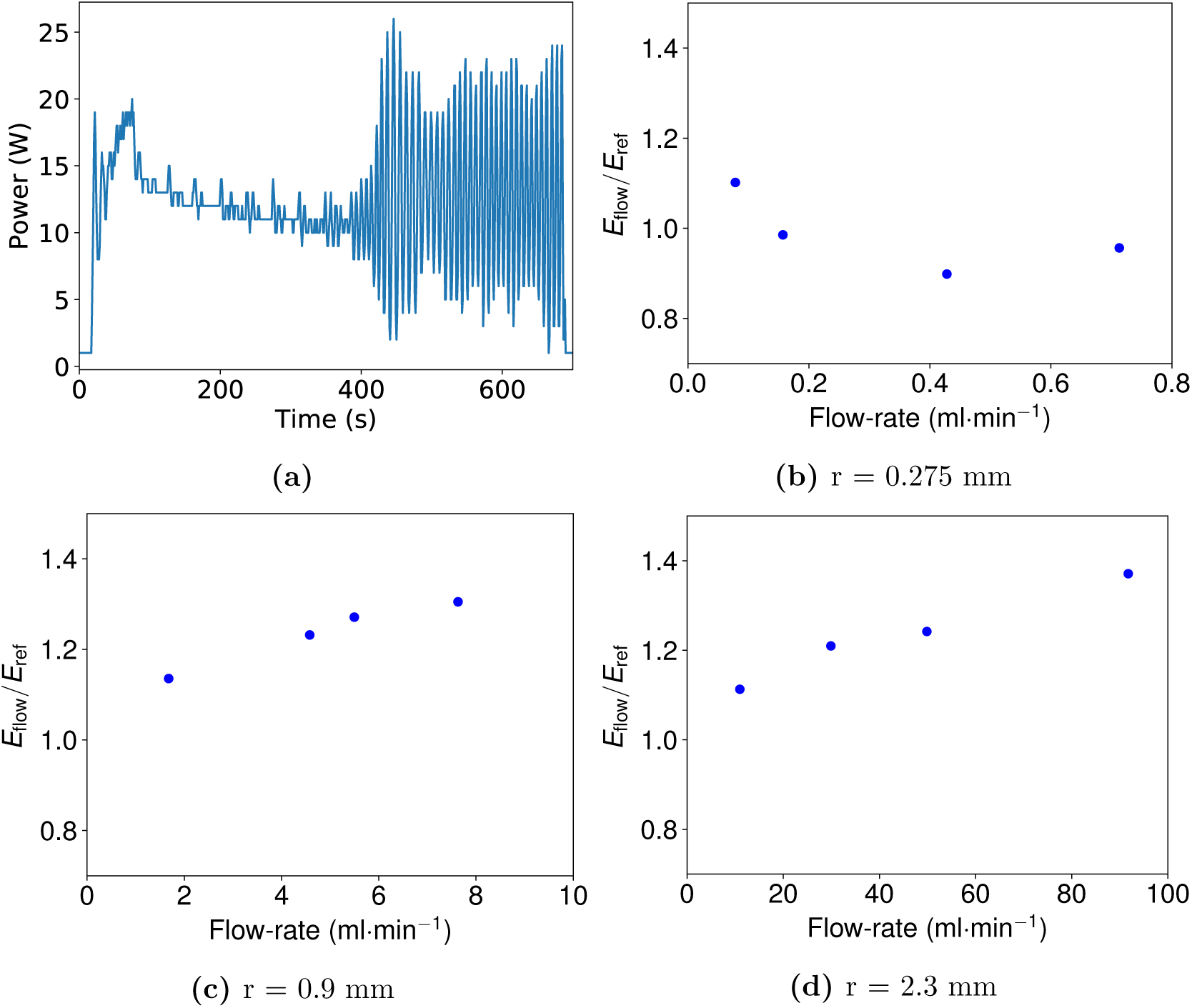
(a) Example power curve that was obtained from the generator logging. (b)-(d) Normalized energy input for the RF ablation experiments with different radii and flow rates.

## 4 Discussion

In this study, ablation experiments were conducted in phantoms with different flow channel radii and flow rates using a temperature-controlled RF generator and a monopolar RF electrode. The experimental results showed an influence of the flow channel radius on the ablation outcomes. Our finding that the measured T> 70°C zone areas did not always decrease with flow rate, but increased for the largest flow channel radius, seemed surprising at first sight, as larger flow rates should result in more heat transport away from the T> 70°C zone and hence in smaller T> 70°C areas. This difference in behavior for the different channel radii can be understood when taking into account the temperature-based feedback algorithm of the RF generator. The generator uses the temperature measured by a probe in the needle tip as feedback in the power control algorithm. A blood vessel of small radius probably reduces the ablated area locally, however, this heat loss is not perceived by the temperature sensor in the needle tip. Larger vessels, on the contrary, draw higher amounts of heat away from the ablation site, so that the overall tissue heating is reduced. Consequently, a lower temperature is measured at the needle tip, which the RF generator compensates for with an increased power input. The increased energy delivery into the tissue then leads to an overall increasing T> 70°C zone area, which is however asymmetric with respect to the needle axis. The results underline that it is not always straightforward to predict the resulting ablation zone size by looking at nearby larger vessels before the start of a treatment - at least not in the case of feedback-controlled RF generators which deliver variable power to the tissue depending on an input variable such as the tissue temperature or impedance. Here, a clear advantage of including mathematical model-based treatment planning into ablation treatments becomes apparent. In the clinical application, monitoring the power input over time and predicting the lesion size in real-time during the treatment could help to define an optimal end point of an RFA. It should be emphasized that our finding applies only to RF generators with needle temperature controlled power delivery: e.g., Bitsch at al. found the ablation zones to decrease under perfusion; however, their generator followed a fixed power curve and not a fixed temperature-controlled power input [7].

The polyacrylamide gel phantoms proved suitable for the flow channel experiments, since a flow channel could be molded directly into it without the need for a glass tube. This resembles the in-vivo situation more closely, as glass vessels were reported to locally change the electric fields [18]. In the flow experiments, saline solution was used as a perfusion fluid due to simplified handling in comparison to blood. This means, however, that our experiments were not suitable to investigate heat-induced coagulation effects. The discoloring of the thermochromic ink was used to mark the *T* > 70°C zone, which should not be confounded with the zone of coagulated tissue. The latter one would have a larger extent, as (1) coagulation also happens at lower temperature and (2) depends on the temperature over time, i.e., on the thermal dose [25]. Although the utilized thermochromic ink did not permit to assess the thermal dose delivered to a tissue, it allowed for a comparison of the *T* > 70°C zone to the results of finite volume simulations.

The investigation into the directional effect suggests that lower channel radii are necessary for the occurrence of a directional effect. Of the three channel radii considered in this work, only experiments using the lowest channel radius of 0.275 mm showed a directional effect. For this radius, the directional effects were seen for the physiological flow-rate (given by Murray’s law) and two other flow rates (one lower than physiological and the other higher). This suggests that as flow-rate is increased for a constant channel radius, the directional effect first increases, reaches a maximum, and then decreases. For the case with the largest directional effect, there was a shrinkage of the *T* > 52°C region at the location of saline inflow, affecting the the ablation safety margin locally. On the other hand there was also a local extension of the *T* > 52°C region at the saline outflow, posing a potential threat to sensitive structures. At the same time the *T* > 52°C region does not differ from the reference *T* > 52°C region away from the saline inflow and outflow. A study by Debbaut et. al. estimated that there are about 10 times more meso-scale blood vessels of radius below 0.3 mm in the liver than of radius above 3 mm, which are vulnerable to thermal damage depending on their distance from the ablation needle [26]. This suggests the need for a more nuanced ablation strategy in the vicinity of vessels capable of producing a directional effect, than simply increasing power input or ablation time.

Of the 15 ablation results shown here, four exhibited a notable difference between simulation and experiment. This difference was always in the form of a ‘bump’ in the *T* > 70°C zone. Three different possible causes for this deviation were explored in this work by performing additional simulations (data not shown). Neither a local inhomogeneity in phantom thermal conductivity, current leakage from the insulated ablation needle surface, nor a local contact resistance on the ablation needle active surface produced a similar feature in the simulated results.

Our study simplifies the complicated clinical situation to a large extent and can only be seen as a step towards the implementation of simulation-based treatment planning into RFA interventions. For instance, vessel geometry and distance to the ablation needle are known with high accuracy in our phantom experiments, and can be fed into the simulation framework without the need for medical imaging beforehand. In an in-vivo ablation treatment, it may however be possible to segment the needle position and the larger vessels based on an initial CT scan, on the basis of which treatment planning can still be executed [3]. Importantly, our experiments revealed an influence of the flow conditions on the thermal lesion, and on the power input of the RF generator, which was moreover varying over time because of the temperature-controlled feedback loop. With respect to treatment planning, it would, therefore, be optimal to monitor the delivered power in real-time and to provide an updated prediction of the achieved ablation zone size. This prediction could define an optimal endpoint for an RFA treatment. To give an example, Thanos et al. [14] observed in a study, which involved among others the RITA RF generator, a local tumor recurrence next to blood vessels for which they compensated by setting longer ablation times in a follow-up intervention. In the cases with tumor recurrence, a simulation-based treatment planning would have been of use to predict a patient-specific optimal endpoint directly for the first RFA.

A similar study, which includes the effects of thermal coagulation of blood [27], could be subject to future work. A thermochromic ink that changes its color at a lower temperature would constitute an important improvement for future experiments. Future work could furthermore comprise more than one flow channel and even more complicated vessel structures.

## 5 Conclusion

In this study RFA was performed in thermochromic gel phantoms with a flow channel using a temperature-controlled Angiodynamics RITA 1500X RF generator. The flow channel radius and volume flow rate were varied and flow effects on the *T* > 70°C zone were observed.

First, the directional effects of the flow were observed only for the 0.275 mm channel radius, and the lower three of the four flow-rates considered. With increasing flow-rate, the directional effect was found to first increase, then reach a maximum and finally reduce again. It led to a shrinking of the thermal lesion upstream of the needle active surface and an extension downstream of the needle active surface. This phenomenon has potential implications for the ablative safety margin and damage to sensitive structures.

While the ablated area reduced with increasing flow rate for the smaller two radii, an increase in area and energy input with flow rate was observed for the large flow channel radius. This implies that vessels adjacent to a tumor can have variable effects on the ablation outcome, at least when working with a temperature-controlled RF generator like the RITA system. Moreover, the influence of perfusion on the time-dependent power input into the tissue makes it very complex to predict the treatment outcome even when the distance to an adjacent vessel is known. Our findings show the potential benefits of simulation-assisted, patient-individual RFA treatment planning and guidance for the prediction of RFA outcomes in the presence of blood flow. While simulation-based ablation treatment planning is still in development, this finding is a step towards improved treatment guidance as compared to lesion estimates provided by ablation equipment manufacturers.

## 6 Acknowledgements

The authors would like to thank Ron Steijvers, Philips Healthcare, for helpful discussions in the chemistry lab. This project has received funding from the European Union’s Horizon 2020 research and innovation program under the Marie Skłodowska-Curie (MSCA-ITN EID) grant agreement 642445 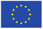.

## References

[1] S. N. Goldberg, G. S. Gazelle, and P. R. Mueller, Thermal ablation therapy for focal malignancy: a unified approach to underlying principles, techniques, and diagnostic imaging guidance, American journal of roentgenology 174, 323–331 (2000).

[2] D. S. Lu, S. S. Raman, P. Limanond, D. Aziz, J. Economou, R. Busuttil, and J. Sayre, Influence of large peritumoral vessels on outcome of radiofrequency ablation of liver tumors, Journal of vascular and interventional radiology 14, 1267–1274 (2003).

[3] S. Payne, R. Flanagan, M. Pollari, T. Alhonnoro, C. Bost, D. O’Neill, T. Peng, and P. Stiegler, Image-based multi-scale modelling and validation of radio-frequency ablation in liver tumours, Philos Trans R Soc Lond A: Math, Phys & Eng Sci 369, 4233–4254 (2011).

[4] M.-H. Park, H. Rhim, Y.-S. Kim, D. Choi, H. K. Lim, and W. J. Lee, Spectrum of CT findings after radiofrequency ablation of hepatic tumors, Radiographics 28, 379–390 (2008).

[5] L. S. Poulou, E. Botsa, I. Thanou, P. D. Ziakas, and L. Thanos, Percutaneous microwave ablation vs radiofrequency ablation in the treatment of hepatocellular carcinoma, World journal of hepatology 7, 1054 (2015).

[6] C. Audigier et al., Comprehensive preclinical evaluation of a multi-physics model of liver tumor radiofrequency ablation, International journal of computer assisted radiology and surgery 12, 1543–1559 (2017).

[7] R. G. Bitsch, M. Düx, T. Helmberger, and A. Lubienski, Effects of vascular perfusion on coagulation size in radiofrequency ablation of ex vivo perfused bovine livers, Investigative radiology 41, 422–427 (2006).

[8] H.-W. Huang, Influence of blood vessel on the thermal lesion formation during radiofrequency ablation for liver tumors, Med Phys 40 (2013).

[9] H. H. Pennes, Analysis of tissue and arterial blood temperatures in the resting human forearm, J of Appl Physiol 1, 93–122 (1948).

[10] A. Nakayama and F. Kuwahara, A general bioheat transfer model based on the theory of porous media, Int J Heat Mass Transf 51, 3190–3199 (2008).

[11] N. Vaidya and M. Baragona and V. Lavezzo and R. Maessen and K. Veroy, Simulation study of the cooling effect of blood vessels and blood coagulation in hepatic radiofrequency ablation, Submitted to Int J Hyperthermia

[12] H. R. Williams, R. S. Trask, P. M. Weaver, and I. P. Bond, Minimum mass vascular networks in multifunctional materials, J R Soc Interface 5, 55–65 (2008).

[13] Y. Kim, W. Lee, H. Rhim, H. Lim, D. Choi, and J. Lee, The Minimal Ablative Margin of Radiofrequency Ablation of Hepatocellular Carcinoma (¿ 2 and ¡ 5 cm) Needed to Prevent Local Tumor Progression: 3D Quantitative Assessment Using CT Image Fusion, American Journal of Roentgenology 195, 758–765 (2010).

[14] L. Thanos, S. Mylona, P. Galani, M. Pomoni, A. Pomoni, and I. Koskinas, Overcoming the heat-sink phenomenon: successful radiofrequency thermal ablation of liver tumors in contact with blood vessels, Diagnostic and Interventional Radiology 14, 51 (2008).

[15] K. Pillai, J. Akhter, T. C. Chua, M. Shehata, N. Alzahrani, I. Al-Alem, and D. L. Morris, Heat sink effect on tumor ablation characteristics as observed in monopolar radiofrequency, bipolar radiofrequency, and microwave, using ex vivo calf liver model, Medicine 94 (2015).

[16] F. G. Poch, C. Rieder, H. Ballhausen, V. Knappe, J.-P. Ritz, O. Gemeinhardt, M. E. Kreis, and K. S. Lehmann, The vascular cooling effect in hepatic multipolar radiofrequency ablation leads to incomplete ablation ex vivo, International Journal of Hyperthermia 32, 749–756 (2016).

[17] C. Welp, S. Siebers, H. Ermert, and J. Werner, Investigation of the influence of blood flow rate on large vessel cooling in hepatic radiofrequency ablation/Untersuchung des Einflusses der Blutflussgeschwindigkeit auf die Gefäßkühlung bei der Radiofrequenzablation von Lebertumoren, Biomedizinische Technik 51, 337–346 (2006).

[18] A. González-Suárez, M. Trujillo, F. Burdío, A. Andaluz, and E. Berjano, Could the heat sink effect of blood flow inside large vessels protect the vessel wall from thermal damage during RF-assisted surgical resection?, Medical physics 41, 083301 (2014).

[19] A. H. Negussie, A. Partanen, A. S. Mikhail, S. Xu, N. Abi-Jaoudeh, S. Maruvada, and B. J. Wood, Thermochromic tissue-mimicking phantom for optimisation of thermal tumour ablation, International Journal of Hyperthermia 32, 239–243 (2016).

[20] A. S. Mikhail, A. H. Negussie, C. Graham, M. Mathew, B. J. Wood, and A. Partanen, Evaluatoin of a tissue-mimicking thermochromic phantom for radiofrequency ablation, Medical Physics 43, 4304–4311 (2016).

[21] A. H. Negussie, A. Partanen, A. S. Mikhail, S. Xu, N. Abi-Jaoudeh, S. Maruvada, and B. J. Wood, Thermochromic tissue-mimicking phantom for optimisation of thermal tumour ablation, International Journal of Hyperthermia 32, 239–243 (2016), PMID: 27099078.

[22] M. H. Sharqawy, J. H. Lienhard, and S. M. Zubair, The thermophysical properties of seawater: A review of existing correlations and data, Desalination and Water Treatment 16, 354–380 (2010).

[23] M. H. Sharqawy, K. G. Nayar, L. D. Banchik, and J. H. Lienhard, Thermophysical properties of seawater: A review and new correlations that include pressure dependence, Desalination 390, 1–24 (2016).

[24] Angiodynamics INC, RITA 1500X USER’S GUIDE AND SERVICE MANUAL (For software version 8.60 and above).

[25] C.-C. R. Chen, M. I. Miga, and R. L. Galloway Jr, Optimizing electrode placement using finite-element models in radiofrequency ablation treatment planning, IEEE Trans Biomed Eng 56, 237–245 (2009).

[26] C. Debbaut, P. Segers, P. Cornillie, C. Casteleyn, M. Dierick, W. Laleman, and D. Monbaliu, Analyzing the human liver vascular architecture by combining vascular corrosion casting and micro-CT scanning: a feasibility study, J Anat 224, 509–517 (2014).

[27] J. K. Barton, D. P. Popok, and J. F. Black, Thermal analysis of blood undergoing laser photocoagulation, IEEE J Sel Top Quantum Electron 7, 936–943 (2001).

